# Comparative specialization of intrinsic cardiac neurons in humans, mice, and pigs

**DOI:** 10.1101/2024.04.04.588174

**Authors:** John D. Tompkins, Donald B. Hoover, Leif A. Havton, Janaki C. Patel, Youngjin Cho, Elizabeth H. Smith, Natalia P. Biscola, Olujimi A. Ajijola, Kalyanam Shivkumar, Jeffrey L. Ardell

**Author notes:** Department of Internal Medicine, Seoul National University Bundang Hospital, Seongnam, Gyeonggi-do, Republic of Korea (Youngjin Cho). **Correspondence:** John D. Tompkins, PhD Department of Medicine (Cardiology) David Geffen School of Medicine University of California Los Angeles, California 90095 https://orcid.org/0000-0001-9496-7930.

## Abstract

Intrinsic cardiac neurons (ICNs) play a crucial role in the proper functioning of the heart; yet a paucity of data pertaining to human ICNs exists. We took a multidisciplinary approach to complete a detailed cellular comparison of the structure and function of ICNs from mice, pigs, and humans. Immunohistochemistry of whole and sectioned ganglia, transmission electron microscopy, intracellular microelectrode recording and dye filling for quantitative morphometry were used to define the neurophysiology, histochemistry, and ultrastructure of these cells across species. The densely packed, smaller ICNs of mouse lacked dendrites, formed axosomatic connections, and had high synaptic efficacy constituting an obligatory synapse. At Pig ICNs, a convergence of subthreshold cholinergic inputs onto extensive dendritic arbors supported greater summation and integration of synaptic input. Human ICNs were tonically firing, with synaptic stimulation evoking large suprathreshold excitatory postsynaptic potentials like mouse, and subthreshold potentials like pig. Ultrastructural examination of synaptic terminals revealed conserved architecture, yet small clear vesicles (SCVs) were larger in pigs and humans. The presence and localization of ganglionic neuropeptides was distinct, with abundant VIP observed in human but not pig or mouse ganglia, and little SP or CGRP in pig ganglia. Action potential waveforms were similar, but human ICNs had larger after-hyperpolarizations. Intrinsic excitability differed; 93% of human cells were tonic, all pig neurons were phasic, and both phasic and tonic phenotypes were observed in mouse. In combination, this publicly accessible, multimodal atlas of ICNs from mice, pigs, and humans identifies similarities and differences in the evolution of ICNs.

## Introduction

The network of nerves on the heart’s surface is essential to the control of cardiac function. At the heart’s hilum, extrinsic nerves enter and splay across the epicardium, intermingling with neurons within epicardial ganglia, forming collectively an ‘intrinsic cardiac nervous system’ (ICNS)^1–3^. This neural network spans the posterior atria and ventricles, and contains nerve fibers from centrally located preganglionic neurons, postganglionic sympathetic nerves, sensory fibers from paravertebral and vagal ganglia, as well as the postganglionic neurons residing on the heart^4–6^. Functional interconnection of the ganglionic neurons is observed; a network interaction that may be subserved by local circuit neurons^7,8^. The epicardial neurons in turn project to cells and tissues of the heart to regulate cellular function via paracrine signaling. This network constitutes the ultimate phase of cardiac autonomic control, encompassing neural components that provide final modulation of regional cardiac function^5,9–11^. Activity within the network modulates the initiation of each heartbeat at the sinoatrial node (SAN), the rate of impulse transmission through the atrioventricular (AV) node, and the amplitude of calcium transients within cardiac myocytes and vascular smooth muscle cells^12^. This modulation maintains a life-sustaining cardiac output in alignment with both organ-specific homeostatic demands and the overall needs of the organism.

Dysfunction in this regulatory neural network, whether initiated by disease or injury, is detrimental to cardiac electrical and mechanical function and increases the risk for sudden cardiac death^13–17^. Novel clinical approaches have arisen to selectively influence organ function by manipulation of the peripheral nerves innervating organs^18–20^. The epicardial nerves of the heart are recognized as targets of neuromodulation strategies implemented by selective electrical, chemical, and/or genetic manipulation^20,21^.

An understanding of the underlying cell structure and diversity among species is essential for developing an informed approach for neuromodulation strategies targeting epicardial neurons. Here, we present a structural and functional atlas of intrinsic cardiac neurons (ICNs) at macro-, meso- and nanoscales from three species: mice, pigs, and humans. Our focus was on clusters of epicardial neurons within the right atrial ganglionated plexus (RAGP), which contain neurons essential to the regulation of heart rate at the SAN^22–24^. Through our investigation, we identified both conserved and derived attributes of these cells among species. Adaptations in synaptic function, neuronal morphology, and neurochemistry revealed specificities in ICN structure and function across mammals, highlighting divergent specifications of human ICNs from those of mice and pigs. This study represents one of the first efforts to characterize human ICN membrane physiology. The research was made possible through collaboration among a multidisciplinary team and funded from the National Institutes of Health, specifically the Common Fund Program for Stimulating Peripheral Activity to Relieve Conditions (SPARC). Portions of this work were previously shared in poster and abstract form ^25,26^.

## Methods

Animal research was conducted in accordance with United States Federal Regulations as set forth in the Animal Welfare Act, the 2011 Guide for the Care and Use of Laboratory Animals (1), the Public Health Service Policy for the Humane Care and Use of Laboratory Animals, and the policies and procedures established by the University of California - Los Angeles (UCLA) Animal Research Committee.

Use of human cardiac tissues was approved by the UCLA Institutional Review Board (IRB). Tissues were procured consistent with organ donation preferences indicated by the donor and/or after signed consent was obtained from family members in compliance with the bioethical laws of the United States of America, the Uniform Anatomical Gift Act (2006), as well as UCLA policies and procedures for use of human tissue.

### Isolation of epicardial ganglia

Adult C57BL/6J (Jackson Labs) mice (n = 11; 4 male, 7 female, age 10 ± 2 wks) were anesthetized deeply with isoflurane (5%) and exsanguinated. The thoracic cavity, including ribs, heart, and lungs, was removed and placed in ice cold physiologic salt solution (PSS) containing (in mM): 121 NaCl, 5.9 KCl, 1.2 NaH_2_PO_4_, 1.2 MgCl_2_, 25 NaHCO_3_, 2 CaCl_2_, 8 D-glucose; pH 7.4 maintained by 95% O_2_ - 5% CO_2_ aeration. The heart was excised from the thorax and observed beneath a stereomicroscope to isolate epicardial ganglia by fine dissection. Connective tissue enveloping isolated ganglia, which were devoid of contracting myocardium, was pinned to the SylGard (Dow Corning) floor of a glass bottom petri dish.

Adult Yucatan minipigs (Premier Biosource; n = 11; 5 castrated male, 6 female; age 6 to 8 mo) were sedated with intramuscular telazol (4–6 mg/kg), intubated, and mechanically ventilated. General anesthesia was maintained with inhaled isoflurane (1.5–2.5%) and intravenous boluses of fentanyl (total: 10–30 µg/kg) during surgical preparation by way of median sternotomy to expose the heart. Major vessels were clamped, the heart was removed, rinsed in PSS, and the dorsal right atrium, at the junction of the superior vena cava (SVC) and the inferior vena cava (IVC), was dissected for subsequent dissection of epicardial ganglia beneath a stereomicroscope.

Human hearts (n = 17; 12 male, 5 female; aged 40 to 72yrs) were procured from deceased organ donors following a formal declaration of brain death. Hearts received were designated for biomedical research after being declined for cardiac transplantation. Deidentified organ donor demographic data is provided in *Supplementary Table 1*. The hearts were transported in ice cold crystalloid cardioplegia (University of Wisconsin solution) and were received within six hours after aortic cross clamp. Upon receipt, a portion of the posterior right atrium containing the RAGP (Figure 1C_1_) was immediately excised and the epicardial ganglia were isolated under a stereomicroscope by fine dissection. The isolated ganglia were pinned to the SylGard floor of a glass bottom petri dishes using methods identical to those for pig and mouse tissue.

**Figure 1.**
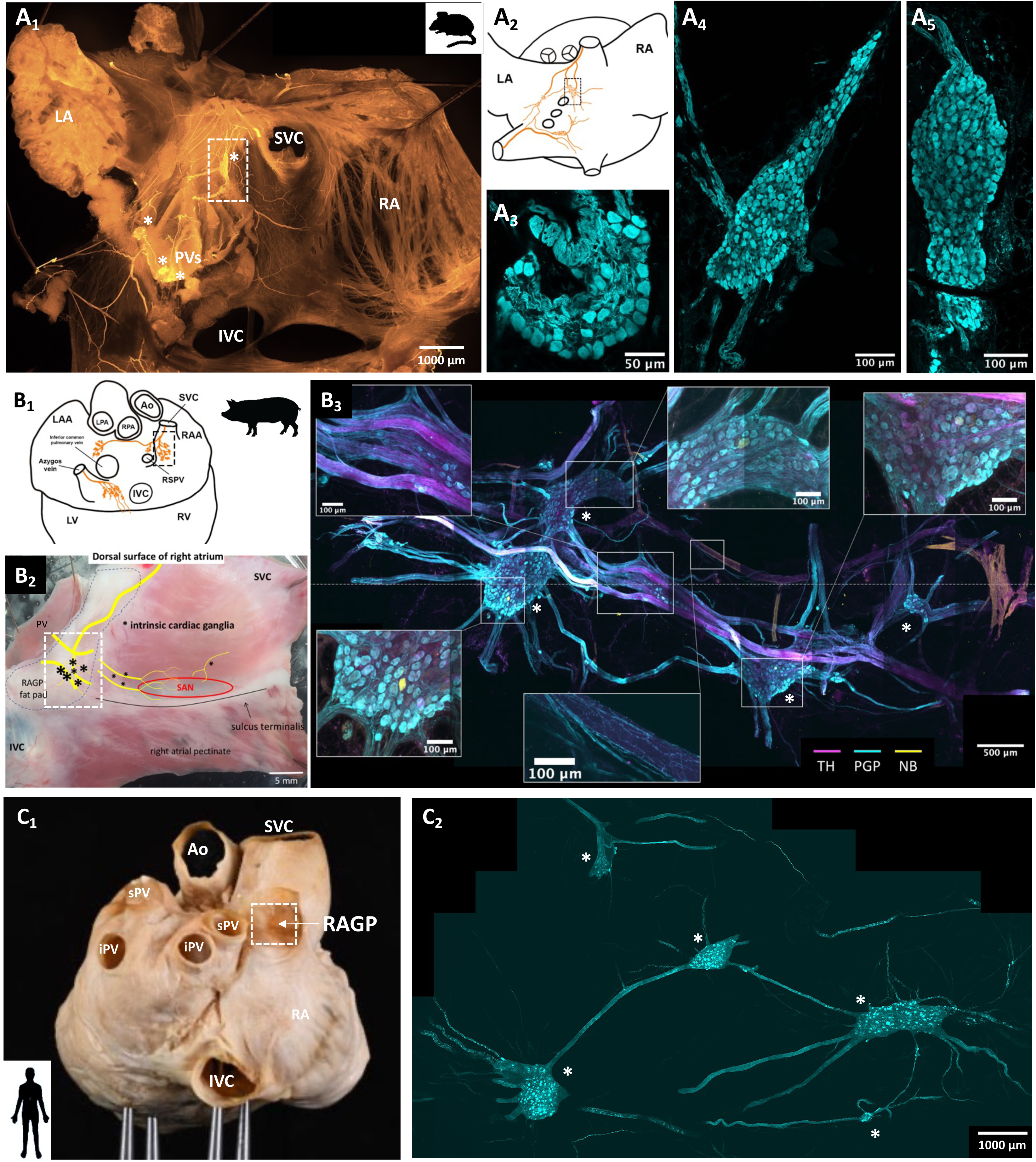
The right atrial ganglionated plexus (RAGP) of mice, pigs and humans. (A_1_) Fluorescent immunolabeling of PGP9.5 in fixed tissue shows intrinsic and extrinsic nerve fibers, and epicardial ganglia (*) on the dorsal surface of the mouse atria. Abbreviations: left atrium (LA); right atrium (RA); inferior vena cava (IVC); superior vena cava (SVC); pulmonary veins (PVs). (A_2_) Ganglion locations illustrated on mouse heart drawing. Boxed region shows location of mouse RAGP. (A_3-5_) Three mouse RAGPs, isolated from boxed regions identified in A_1&2_, showing immunofluorescence for PGP9.5. (B_1_) Cartoon illustrates location of epicardial ganglia on minipig dorsal atria. (B_2_) Photo of right atrial dorsal epicardium indicating location of RAGP identified by boxed regions shown in B_1_&B_2_. Approximate location of ganglia (*), nerves (yellow lines), and the sinoatrial node (red oval) are shown. (B_3_) An RAGP isolated from the area shown in B_2_. Immunofluorescence to PGP9.5, tyrosine hydroxylase (TH), and injected neurobiotin (NB) are shown. Insets show greater magnification of identified areas. (C_1_) Photo of a human heart after perfusion fixation identifies location of human RAGP isolation. Abbreviations: superior vena cava (SVC); inferior vena cava (IVC); inferior pulmonary vein (iPV); superior pulmonary vein (sPV); right atrium (RA); aorta (Ao). (C_2_) Photomicrograph of an RAGP isolated from a human heart showing immunofluorescence to PGP9.5. Five epicardial ganglia (*) connected by interganglionic nerves are shown.

Isolated ganglion preparations were transferred to the stage of an upright compound microscope (Zeiss AxioExaminer) equipped with differential interference contrast optics, an Axiocam camera, and a 5X air and a 40X water-immersion objective. Ganglia were superfused (6-7 ml/min) continuously with PSS maintained at 34 ± 2 °C.

### Intracellular microelectrode recording

Visually identified neurons were impaled with borosilicate glass microelectrodes filled with 2% neurobiotin (Vector Labs, Burlingame, CA) in 2.0 M KCl. Electrode tip resistances measured between 60 and 120 MΩ. Recordings of membrane potential were made with a Multiclamp 700B amplifier (Molecular Devices, Sunnyvale, CA) in current clamp configuration. Data was acquired using a Digidata 1550B (Molecular Devices) analog-to-digital converter. The amplifier and data acquisition were controlled with pCLAMP software (Version 10, Molecular Devices).

Depolarizing and hyperpolarizing currents were injected through the recording electrode to determine responsiveness of the neuronal membrane. Hyperpolarizing current steps (500 ms duration; 50-500 pA) measured whole cell input resistance and membrane time constant (tau). Steady-state membrane potential changes in response to both hyperpolarizing and depolarizing currents were assessed. Depolarizing current steps (500 ms duration) of increasing intensity were used to measure membrane excitability. Excitability curves were produced by plotting the number of action potentials elicited by the range of depolarizing currents^27^. While the data denotes a continuous variable (spikes/time), they are presented as spike counts for both clarity and simplicity. Cells were classified as either phasic (firing ≤4 spikes) or tonic (firing >4 spikes continuously) based on the number of action potentials evoked by long-duration, supramaximal depolarizing currents (500 ms). The amplitude and duration of the action potential upstroke were measured from spontaneous or nerve evoked action potentials. The amplitude and duration of the after-hyperpolarizing potential was measured from action potentials evoked by brief intracellular current injection (5 ms duration). Cumulatively, the following membrane properties were assessed: resting membrane potential, rheobase, whole cell input resistance, tau, AP 1/2 width, AP duration, AHP amplitude, 2/3^rds^ AHP duration.

Synaptic potentials were evoked within intact ganglion preparations by focal stimulation of interganglionic nerves using extracellular, concentric bipolar electrodes (FHC; Bowdoin, ME). Graded shocks (50-800 uA; 100 µs) were delivered by an AMPI Master 8 and IsoFlex optical Stimulus Isolation Unit. Stimulus recruitment curves were generated by plotting the latency of the evoked potential against the stimulus current. Five to 20 stimuli were delivered at each stimulus intensity, at an interval of three seconds between stimuli. Latency was calculated from the onset of the command pulse to the upstroke of the evoked potential. Trains of stimuli were given at 5, 10, or 20 Hz to assess spike following frequency at the post-synaptic neuron. The amplitudes of spontaneous excitatory post-synaptic potentials (sEPSPs) were quantified using MiniAnalysis software (Synaptosoft, Inc.). The following synaptic properties were assessed: latency, spike following frequency, sEPSP amplitude.

### Tissue fixation and IHC

After intracellular recording, ganglia were fixed by overnight immersion in 4% paraformaldehyde at 4 °C. The fixed tissue was rinsed in phosphate buffered saline (PBS) and stored in PBS + 0.02% sodium azide at 4 °C until subsequent immunohistochemical processing. For immunostaining of ganglion whole mounts, tissue was blocked in 0.01M PBS, 0.02% sodium azide, 0.1% Triton X-100, and horse serum for four hours at room temperature with agitation. The tissue was then incubated in primary antibodies, in a solution of 0.01M PBS, 0.02% sodium azide, and 0.1% Triton X-100, with agitation over two nights. Tissues were rinsed in a solution of 0.01M PBS, 0.02% sodium azide (1hr rinse, 3 rinses) and incubated in secondary antibodies, diluted in 0.01M PBS with 0.1% Triton X-100 and 0.02% sodium azide, over two nights at room temperature with agitation. Stained tissue received a final rinse in 0.01M PBS, 0.02% sodium azide (1hr rinse, 3 rinses) and was mounted on glass slides in anti-fade media (Citifluor, Electron Microscopy Sciences) and coverslipped. The source and concentration of primary and secondary antibodies is given in Table 1.

**Table 1.**
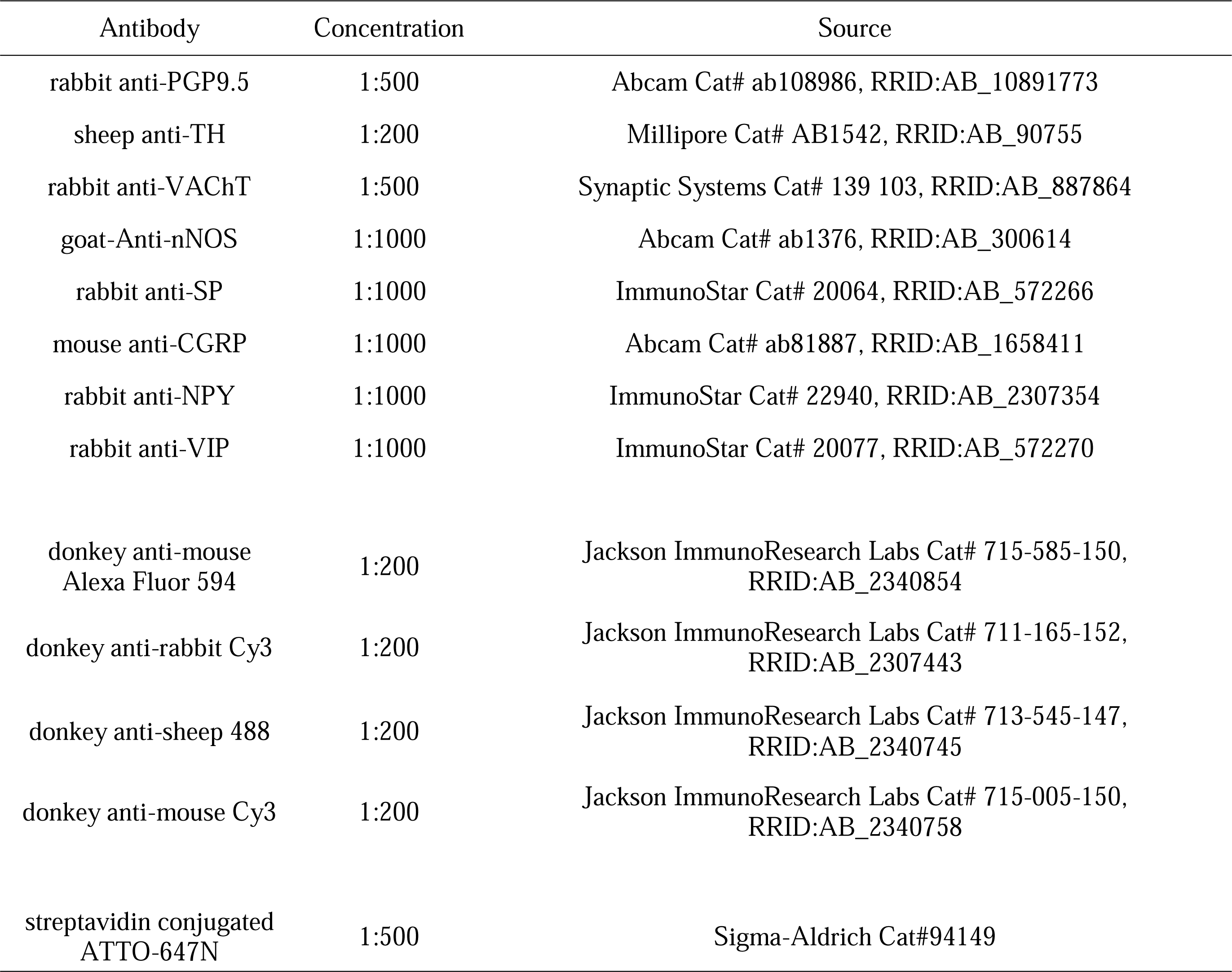
List of antibodies, sources, and concentrations used.

### IHC of sectioned tissue

Tissue blocks of the RAGP fat pad were frozen on dry ice and stored at −80 °C until sectioning. Blocks were attached to specimen plates, cut at −20 to −25°C using a Leica CM3050S cryostat (Leica Microsystems Inc.), and 30 μm thick sections collected on charged slides. Tissues were sectioned in a plane parallel to the epicardial surface, and sections were collected in a sequence that yielded eight sets of sections that each spanned the entire thickness of the specimen. Each set of tissue sections was stored at −20 °C until further processing. Slide-mounted tissue sections were immunostained at room temperature for markers using standard methods of fluorescence immunohistochemistry, as described previously^28^. Briefly, slides were rinsed with 0.01 mol/L PBS (pH 7.3), incubated for 10 minutes in 0.01 mol/L PBS containing 0.4% Triton X-100 and 0.5% bovine serum albumin (BSA), and blocked for 2 hours in 0.01 mol/L PBS containing 1% BSA, 0.4% Triton X-100, and 10% normal donkey serum (Jackson ImmunoResearch Laboratories). Each section was then incubated overnight in blocking buffer containing one or more of the primary antibodies listed in Table 1. Sections were washed again and incubated in blocking buffer before application of secondary antibodies. Species-specific donkey secondary antibodies conjugated to Alexa Fluor 488, 555, 594 or 647 (Jackson ImmunoResearch Laboratories) were applied at a 1:200 dilution in 0.01 mol/L PBS containing 0.4% Triton X-100 and 1% BSA, and sections were incubated for 2 hours before final washing in 0.01 mol/L PBS. After final washes with 0.01 mol/L PBS, cover glasses were applied with Citifluor (Ted Pella, Inc.) or SlowFade Gold antifade reagent (Life Technologies Corporation). Specific staining did not occur in negative control sections processed without the addition of the primary antibodies.

Slides were viewed under fluorescence illumination with an Olympus BX41 microscope equipped with an Olympus DP74 digital camera and cellSens software (Olympus America Inc., Center Valley, PA, RRID:SCR_016238). After localizing ganglia within sections, they were evaluated with a Leica TCS SP8 Confocal Microscope with 10x, 20x and 40x objective lenses (Leica Microsystems Inc.). Confocal images were collected at a resolution of 1024 x 1024 using 488 and 552nm laser lines. Stacks spanned tissue thicknesses of 25-33µm unless otherwise noted. Figures were created using maximum intensity projection (MIP) images for individual channels and merged images.

Whole ganglia were imaged with a confocal microscope (Zeiss LSM880) by tile-scanning in X, Y, and Z planes using a 63X objective (Zeiss Plan-Apo, 1.4NA). Z-stack images were acquired at step sizes consistent with Nyquist sampling in relation to the numerical aperture of the objective. Three-dimensional scans of whole ganglion neuroanatomy were acquired with 10X objectives. Images were quantified using ImageJ to measure neuronal cross-sectional areas (measured at Z-plane containing nucleus), and total cell counts per ganglia were quantified with ImageJ (FIJI distribution).

### Quantification of cellular morphology

Neurobiotin filled cells were serially imaged to visualize cell structure. For cases in which a neurobiotin labeled soma could not be located, or there were not enough dendrites labeled in the volume, the cell was omitted from further analysis. Neuronal structures were segmented, and quantified, using Neurolucida 360 and Neurolucida Explorer (MBF Bioscience). The file format of the traced structures was converted from .dat to .swc using HBP Neuron Morphology Viewer^29^. SWC files were further quantified using the ImageJ plugin SNT^30^. Quantified values include the following: cell soma and dendrite area in X, Y and Z planes, soma volume, number of dendrite stems (originating branches from soma), number of dendrite bifurcations, number of dendrite branches, number of dendrite tips, total dendrite length, mean dendrite length, total dendrite surface area, total dendrite volume.

### Transmission electron microscopy

Samples of whole-mount right atrial ganglion plexi (RAGPs) intended for electron microscopy (EM) analysis were carefully isolated from the heart using a dissection microscope and then submerged in a solution containing 2% paraformaldehyde (PFA) and 2.5% glutaraldehyde overnight at 4°C in a 0.12 mol/L Millonigs buffer (MB). After thorough washing with Millonigs buffer and double-distilled water (ddH_2_O), the tissues underwent fixation in a 1% osmium solution diluted in ddH_2_O, followed by a series of ethanol (EtOH) and propylene oxide dehydration steps. Subsequently, the tissues were embedded in a plastic resin (Epon, Ted Pella). Semi-thin cross-sections (0.5 μm thick) were prepared and stained with a 1% toluidine blue solution diluted in ddH_2_O to enable a general overview of RAGP organization under a light microscope (Nikon Eclipse E600 microscope equipped with a Nikon DS-Fi3 camera). Ultra-thin sections (70 nm thickness) were then mounted onto formvar-coated grids (Ted Pella), stained with uranyl acetate and lead citrate for contrast enhancement, and examined in a Tecnai G2 Spirit Twin transmission electron microscope (FEI, ThermoFisher Scientific) operating at 80LkV. Detailed characterization of epicardial ganglion organization was performed, and images were captured using a Gatan Orius SC 1000B digital camera (Gatan) and SerialEM software^31^.

### Statistical analysis

Statistical analysis was performed using GraphPad Prism statistical software (version 9.2; La Jolla, CA). Data are presented as meanL±LSD. Means of two independent groups were compared with an unpaired t test. Categorical data arranged into contingency tables were analyzed using Fisher’s exact test. Grouped data with two categorical variables and one continuous dependent variable were compared by two-way ANOVA and Šidák’s multiple comparisons test. Means between three groups were compared with one-way ANOVA and Tukey’s post hoc analysis. For datasets with missing values a mixed-effects analysis model was used. Values were considered statistically significant at *P* ≤ 0.05.

### Data Availability

Data associated with this study were collected as part of the Stimulating Peripheral Activity to Relieve Conditions (SPARC) program and are available through the SPARC Portal (https://sparc.science/) (RRID: SCR_017041) under a CC-BY 4.0 license.

## Results

### Macroscopic neuroanatomy of epicardial neurons

In mice, epicardial ganglia are found on the epicardial surface beneath a layer of connective tissue on the dorsal atrial surface near the pulmonary veins (Fig 1A). Approximately 4-7 large ganglia were identified following fixation and incubation with an antibody to the pan-neuronal protein PGP9.5 (Fig 1A). Counted ganglia (n = 6) contained 130 ± 116 neurons, with a mean ganglion area of 54 ± 45 mm^2^, and neuron area of 267 ± 73 µm^2^, giving a total neuronal density of 61 ± 37 neurons per 100 µm^2^. From this data, we estimate the population of mouse ICNs to be between 500 and 1000 total neurons. This is the first quantification of ganglion cell numbers using whole isolated ganglia and three-dimensional imaging with confocal microscopy. Representative quantified ganglia are shown in Figure 1A. For single cell electrophysiology and morphology, we focused on neurons within the ganglion located at the merger of the superior and inferior vena cava, as indicated in Figure 1A.

In adult pig and human cardiac tissues, epicardial ganglia were more challenging to visualize given a thick layer of enveloping subepicardial fat. We focused on clusters of epicardial ganglia referred to commonly as the RAGP. The localization of the RAGP in pig and human hearts is shown in Figures 1B and 1C. Neurons from these ganglia primarily project to the right atrial myocardium and sinoatrial node^22–24^. Macroscopically, in pig and human, the RAGP appeared as an interconnecting web of nerve bundles with interspersed ganglia of varying size, consistent with the nomenclature of a ‘ganglionated plexus’ (Fig 1B_3_ & C_2_). This neuroanatomy, while conserved between animals, showed significant interspecies, and more subtlety, inter-animal variability. The areas of individual ganglia, 168 ± 200 mm^2^ in pig (n = 30) and 162 ± 28 mm^2^ in human (n = 19), and cross-sectional area of the neuronal soma (1133 ± 27 µm^2^ vs 1124 ± 45 µm^2^) were similar between pig (n = 298) and human (n = 187). The neuronal density between the pig (n = 4) and human (n = 4) were also similar (6.7 ± 0.5 neurons/100 µm^2^ vs 4.0 ± 0.4 neurons/100 µm^2^). Despite a similar mean ganglion size, the density of the smaller mouse neurons was greater than pig (P = 0.0034) or human (P = 0.0054).

### Immunohistochemical observations of neurotransmitter diversity

Cellular neurochemical expression was probed with rigorously validated antibodies (Table 1), in both ganglion sections and whole mounts. In all three species, intense punctate labeling of the vesicular acetylcholine transporter (VAChT) was observed surrounding most neuronal perikarya, supporting a ubiquitous preganglionic cholinergic innervation to these cells. Intracellular VAChT+ labeling was observed in most, but not all cells (Fig 2), indicative of a large population of post-ganglionic cholinergic neurons in epicardial ganglia from mice, pigs and humans.

**Figure 2.**
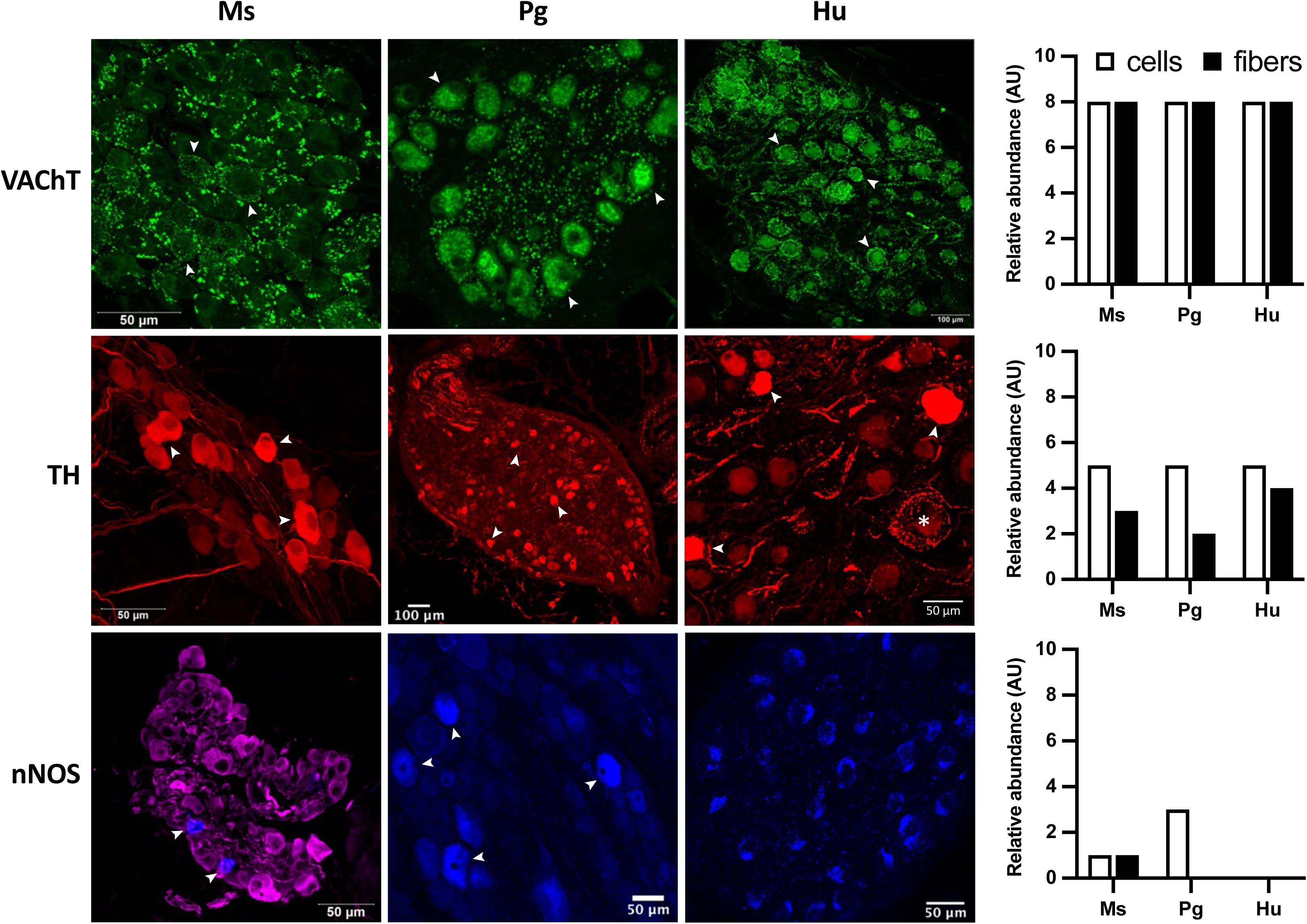
Fluorescently immunolabeled ganglia from mice, pigs and humans showing presence and localization of cholinergic, adrenergic and nitrergic proteins. Confocal photomicrographs of sectioned and whole-mount ganglia show representative stainings for the indicated proteins (vesicular acetylcholine transporter (VAChT); tyrosine hydroxylase (TH); neuronal nitric oxide synthase (nNOS)). Bar graphs show relative abundance of staining within either neuronal cell bodies or nerve fibers of each species after review of multiple images from several animals. Top panel arrows indicate pericellular, cholinergic varicosities and VAChT+ cell bodies in each species. Middle panel arrows indicate TH+ neurons. The asterisk in the human RAGP section indicates a cell with pericellular localization of TH+ nerves. Arrows in bottom panels indicate nNOS+ cells. Note that the staining observed in the human tissue was autofluorescence from lipofuscin. No nNOS+ cells were identified in human ganglia.

Immunolabeling with an antibody to tyrosine hydroxylase (TH), a marker for noradrenergic neurons, revealed moderate to strong staining of the neuronal cytoplasm for some ICNs (Fig 2). While cell bodies were strongly positive for TH in each species, only within human ganglia did we clearly observe TH-immunoreactive (IR) nerve terminals encircling the postganglionic neurons (Fig 2).

In addition to adrenergic and cholinergic cells, nitrergic neurons have also been identified within intracardiac ganglia^5^. Using an antibody to the neuronal isoform of nitric oxide synthase (nNOS)(Table 1), sparse cytoplasmic immunolabeling of nNOS was observed in several neurons from mouse and pig, but not human (Fig 2). No intraganglionic nNOS-IR fibers were observed in sections or whole mounts of cardiac ganglia from mice, pigs or humans.

Peptidergic neuromodulators are also present in peripheral ganglia of multiple species, including human^32^. Immunostaining for the sensory neuropeptides substance P (SP) and calcitonin gene-related peptide (CGRP) exposed striking differences across species. In mice, SP and CGRP are colocalized within nerve fibers surrounding neurons as well as within intracardiac nerve bundles; however, no CGRP, or SP-IR cell bodies were identified (Fig 3). Pig intracardiac ganglia showed little to no staining for CGRP or SP containing perineuronal fibers or cell bodies. We only identified a few dually positive fibers within the intracardiac nerve bundles. Human intracardiac ganglia showed colocalization of CGRP and SP, like mouse, and displayed a dense meshwork of perineuronal fibers, also as observed in mice. A couple of human ICNs appeared to exhibit immunoreactivity (IR) to these peptides, though a definitive confirmation of sensory neurons within intracardiac ganglia is lacking.

**Figure 3.**
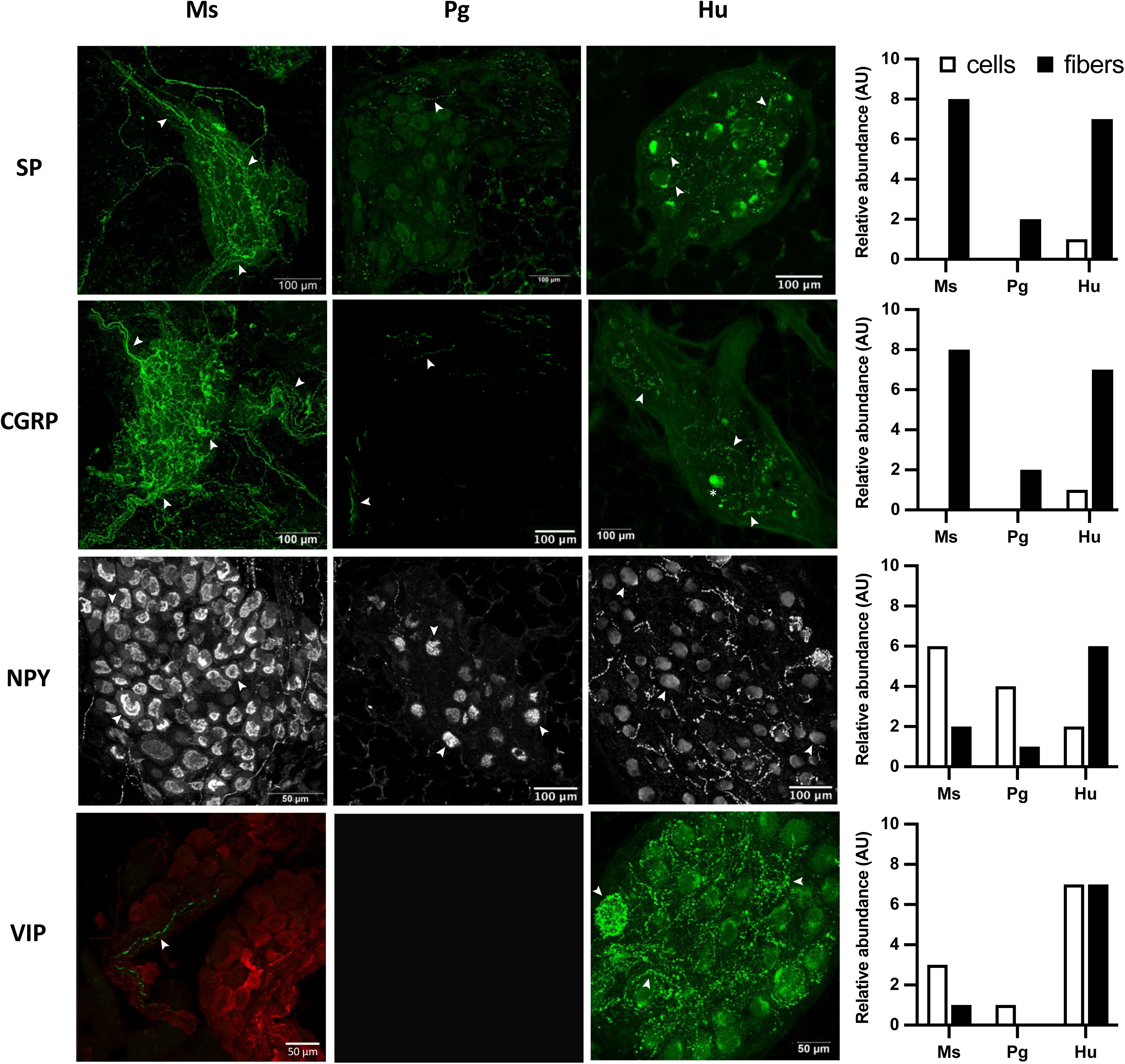
Immunolabeled ganglia from mice, pigs and humans show presence and localization of the indicated neuropeptides. Confocal photomicrographs of sectioned and whole-mount ganglia show representative stainings for the indicated proteins (substance P (SP); calcitonin gene related peptide (CGRP); neuropeptide Y (NPY); vasoactive intestinal polypeptide (VIP)). Right panels show relative abundance of staining between species in each row after review of multiple images from several animals. White arrows in top two row indicated representative SP+ or CGRP+ nerve fibers. The asterisks in human ganglion section of second row indicates a putative CGRP+ neuron. Note apparent staining in cytoplasm of human neurons (top row) was autofluorescence from lipofuscin. Arrows in third row show NPY+ neuronal cell bodies. Arrows in bottom row indicate VIP+ intraganglionic nerve fibers.

Neuropeptide Y (NPY) and vasoactive intestinal polypeptide (VIP) are two peptidergic co-transmitters commonly identified within sympathetic and parasympathetic neurons, respectively^33–35^. In the mouse, NPY-IR was observed in cell bodies, a few fibers, and was often colocalized with TH in nerve fibers surrounding blood vessels (Fig 3). In the pig, few NPY-IR nerve fibers were identified. NPY was principally localized to cell bodies and some co-labeling with TH was observed (Fig 3). In human, a greater abundance of NPY-IR nerve fibers surrounding postganglionic neuronal somata was identified. Perineuronal VIP-IR nerve fibers were also significantly more abundant in human ganglia compared to pig or mouse. Within pig RAGP, only a few VIP-IR cell bodies were identified. In mouse, a few varicose VIP-IR fibers were observed.

Importantly, immunoreactivity for the peptides showed variability across ganglia. The above observations were inferred from imaging of 18 sections and 2 whole mounts from 3 mice, 27 sections and 2 whole mounts from 5 pigs, and 32 sections and 2 whole mounts from 4 humans.

### Passive and active membrane properties

Intracellular microelectrode recordings from intact ganglion preparations were used to assess basic membrane properties of ICNs from each species (Fig 4). Recordings were obtained in identical extracellular solutions made at 34 ± 2 °C. Representative action potentials from each species are overlaid in Fig 4D and show similar initial rates of depolarization and repolarization. The mouse cells rested at less negative potentials (−48 ± 8 mV), relative to pig (−58 ± 8 mV) or human (−54 ± 8 mV)(Fig 4E), and correspondingly had shorter action potentials (Fig 4F). Action potential half-widths were similar between groups (Fig 4G). The overlaid after-hyperpolarizing (AHP) potentials illustrate the markedly larger AHP of human ICNs, which differed in amplitude but not duration between species (Fig 4H,I,J). The Human AHP amplitudes (−14.5 ± 4 mV) were greater than those of mouse (−9.8 ± 2 mV; P< 0.0001) or pig (−11.3 ± 3; P = 0.0064). The smaller mouse ICNs had higher whole cell input resistance (158 ± 109 MΩ), in comparison to the larger pig (101 ± 58 MΩ) and human (83 ± 36 MΩ) neurons (Fig 4K). The threshold current for action potential general (rheobase) was less in mouse (60 ± 26 pA, n = 20) compared to human (179 ± 70 pA, n = 14; P = 0.0124) or pig (435 ± 161 pA, n = 29; P < 0.0001) neurons. The pig neurons required significantly more current to elicit an action potential compared to human (P <0.0001).

**Figure 4.**
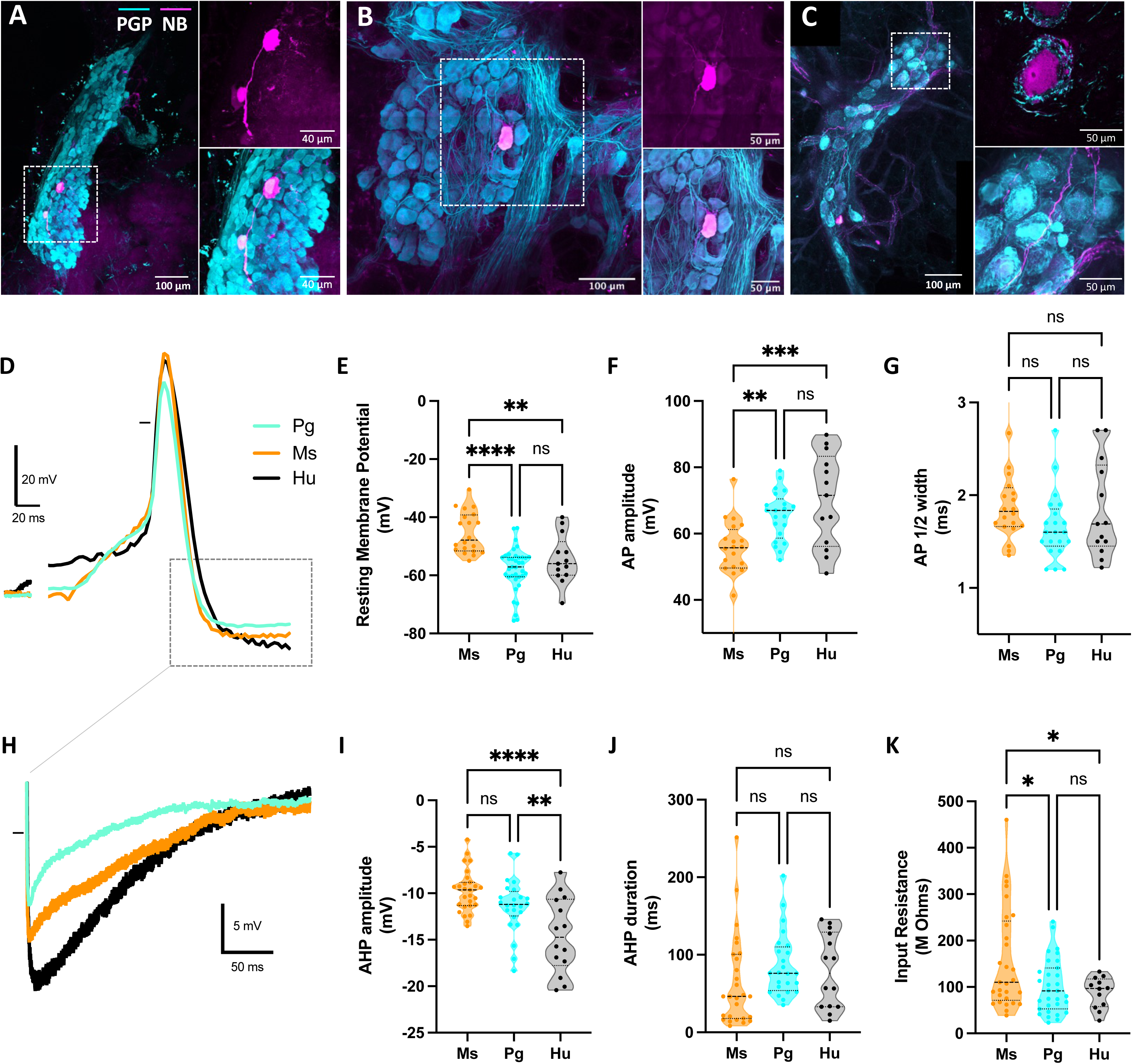
Passive and active membrane properties of ICNs recorded with intracellular microelectrodes. (A-C) Confocal images of right atrial ganglia isolated from a mouse (A), a minipig (B), and a human (C). ICNs were visualized with fluorescently labeled antibodies to PGP9.5 (cyan) and intracellularly injected neurobiotin (magenta). White boxed area in left panels show at higher magnification to right. (D) Overlaid action potentials from a mouse (orange), a pig (cyan), and a human (black) ICN. (E) Resting potentials of mouse neurons were less negative than pig or human ICNs. (F) The amplitude of nerve evoked or spontaneous action potentials was greater at pig and human ICNs. (G) Action potential durations were similar across species. After-hyperpolarizations (AHPs) from the APs shown in (D) are overlaid. (I) The AHPs recorded from human ICNs were greater in amplitude than either mouse or pig AHPs. (J) AHP duration varied greatly across species, with both short and long-lasting AHPs observed. There was not difference in mean amplitude between species. (K) The smaller mouse ICNs had higher input resistance, evidenced by a larger shift in membrane potential in response to hyperpolarizing currents.

Excitability of the post-synaptic membrane was assessed in response to long duration, depolarizing current steps (+50 to +800 pA, 500ms). Autonomic neurons are characterized based on the number of action potentials elicited by depolarizing stimuli^36^. Phasic cells fire short bursts of action potentials in response to long depolarizing current steps, while tonic cells fire steadily throughout the depolarization. In mice, both phasic and tonic neurons were identified (Fig 5A). In pig, no tonic neurons were observed (0/37), and in the human, mostly tonic cells (13/14) were observed (Fig 5B,C). Tonic human ICNs reached a peak firing frequency of 23 spikes/500ms (46 Hz) while mouse cells peaked at twice that frequency (45 spikes/500ms; 90Hz).

**Figure 5.**
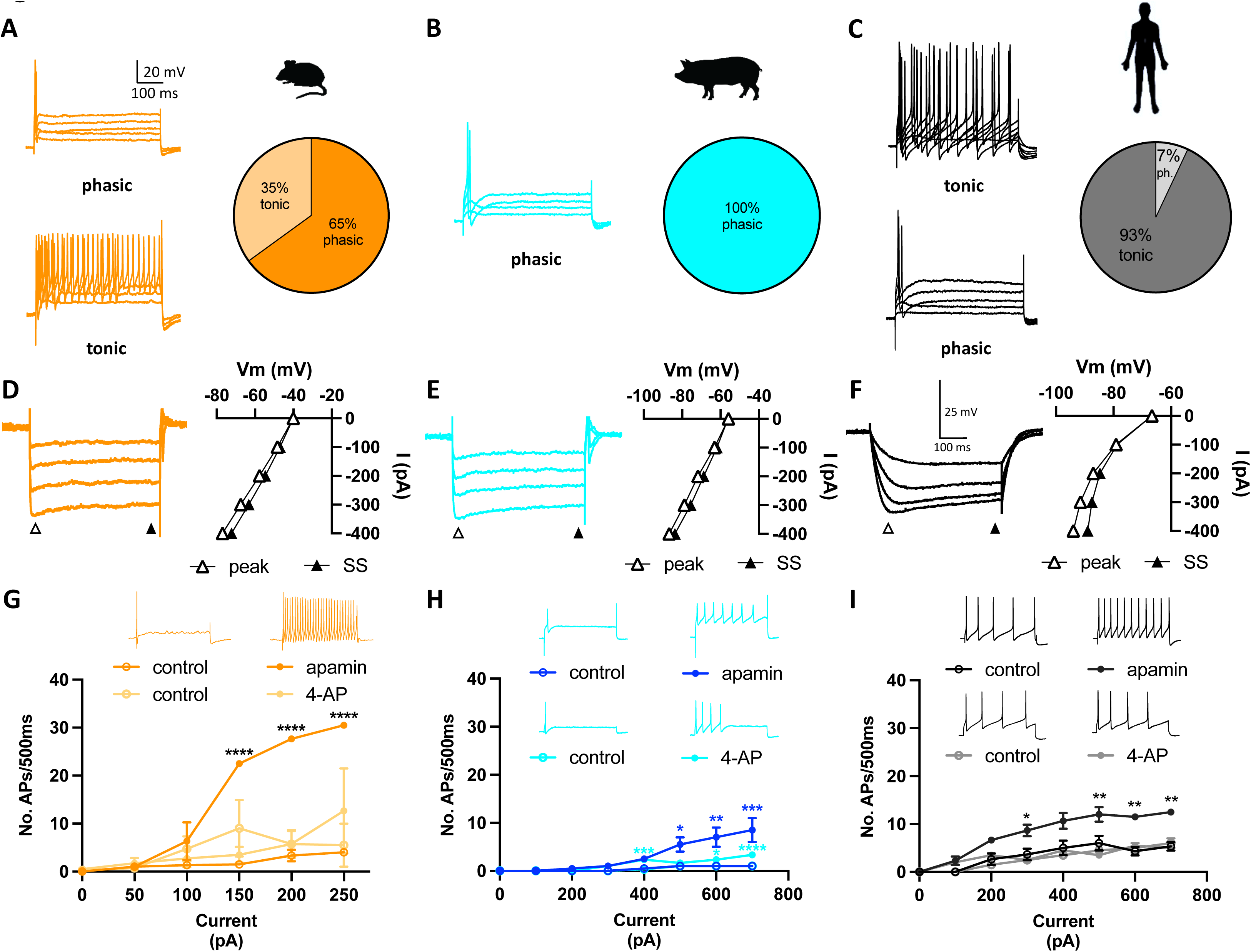
Regulation of membrane excitability varies across species. (A) In mice, phasic and tonic neurons were identified (n = 31). (B) In pig, no tonic cells were observed out of 37 sampled ICNs. (C) In human, all but one cell was tonically firing (n = 14). (D-F) Membrane potential responses to hyperpolarizing current (−100 to −400pA) are shown. The peak (open triangle) and steady-state (closed triangle) change in membrane potential is plotted at right. Triangles indicate where the potential change was measured. Sag in the hyperpolarization, likely due to activation of H-current, was observed in all three species and is shown in representative recordings from each species. (G-I) Pharmacological antagonism of K^+^-currents known to be expressed at ICNs were studied. Inhibition of A-current with 4-AP increased excitability of pig (H) but not human ICNs. 4-AP both increased and decreased excitability in mouse neurons (G). Apamin greatly increased the excitability of mouse (G), pig (H) and human (I) ICNs, causing greatest increase in mouse ICN firing frequency.

The membrane time constant, tau, was calculated from −100pA hyperpolarizing steps. Mouse cells had the shortest charge time (2.4 ± 1 ms, n = 30) relative to human (16.6 ± 13.6 ms, n = 14; P < 0.0001) or pig (9.7 ± 8 ms, n = 33; P = 0.0011). Despite being similar in size, pig neurons charged faster than human cells in response to −100pA steps (P = 0.0178). Larger amplitude, long duration (500ms), hyperpolarizing steps (−100 to −600 pA) were given to assess responsiveness to membrane hyperpolarization. A difference between the instantaneous and steady-state voltage response (Fig 5D-F) indicated the presence of a hyperpolarization-activated non-selective cationic current, known to be mediated by hyperpolarization-activated cyclic nucleotide-gated (HCN) channels^37^, which opposes the hyperpolarization. This change in the hyperpolarization is commonly termed ‘sag’. A ‘sag’ or difference potential of greater than 2 mV was observed in 75% human ICNs, 33% of porcine ICNs, and 85% of mouse ICNs.

Both the A-type voltage-gated potassium channels and small conductance Ca^2+^-activated potassium (SK) channels have been shown to affect intrinsic excitability of mammalian ICNs^38,39^. 4-AP, an inhibitor of A-type channels, increased excitability of pig but not human RAGP ICNs (Fig 5H,I). 4-AP (10 μM), slowed repolarization of the action potential which was measured as an increased action potential duration in mouse, pig and human cells (data not shown). Apamin, an inhibitor of SK channels, significantly increased the number of action potentials elicited with depolarizing current steps in all three species (Fig 5G,H,I). In mouse and pig, phasic neurons were converted to tonically firing ones. After 5 mins, apamin (100 nM) significantly reduced the duration of the AHP in human (130 ± 34 ms before, 51 ± 21 ms after, P = 0.03, n = 3) and mouse (81 ± 20 ms before, 11.5 ± 2 ms after; P = 0.02, n = 3), but not pig ICNs (62 ± 11 ms before, 52 ± 16 ms after; P = 0.2, n = 2).

### Morphologic properties of single ICNs

The segmented cellular morphologies of representative neurons from each species are shown in Figure 6. Mouse neurons (n = 7) were unique from pig or human ICNs in that few or no dendritic projections were observed with light microscopy. In only 1/7 cells were any small dendrites observed. The small diameter neuronal cell bodies projected a single axon which typically, though not always, exited the ganglion (Fig 6A_1-4_). In contrast, the segmented neurons from pig (n = 9) and human (n = 6) exhibited elaborate dendritic arbors consisting of multiple branches projecting from the soma (Fig 6B,C). Typically, these projections extended from the basal membrane, while the apical membrane was smooth. A couple of cells which were deeper within the ganglion appeared to have projections extending from both basal and apical membranes.

**Figure 6.**
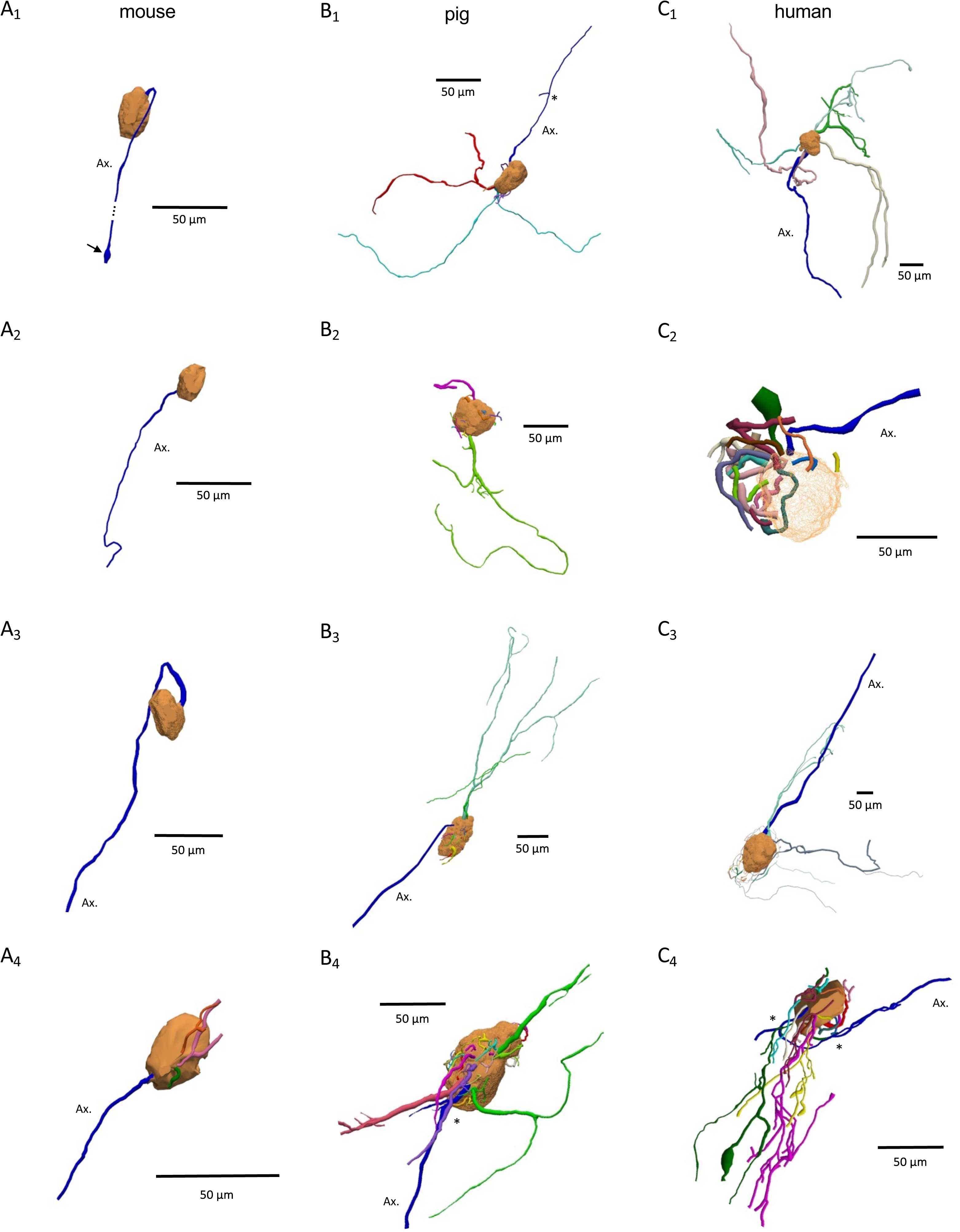
Morphologies of mouse, pig and human ICNs. Segmentation of neurobiotin filled cells reveals diversity of cellular morphologies among species. (A_1-4_) Mouse ICNs have a single axon (Ax.) and most cells have no dendrites. Arrow in A_1_ indicates presumed axon growth cone. Pig (B_1-4_) and human (C_1-4_) ICNs have complex dendritic arbors and typically a single axon. The neuron in B_2_ did not appear to have an axon. Dendritic branches for human cell in C_2_ stayed very close to cell body. All axons are shown in blue. Cell bodies are light orange. Each dendritic stem and all its branches match in color. Axon bifurcations (*) at pig and human ICNs gave rise to short collateral axon branches.

Axons were identified based on larger cross-sectional diameter of the initial segment, and as the longest projections which exited the ganglion to join a nerve bundle. These projections are labeled ‘Ax’ on the shown reconstructions (Fig 6). Axons appeared to originate near dendrites on the soma. Given that the ganglia were isolated from the whole heart, the eventual targets of the axons could not be determined. Some pig and human ICN axons showed small bifurcations near the soma, giving rise to one or two short collaterals (Fig 6B,C). The 3D confocal images and 3D cellular reconstructions are available online (see *Data Availability* statement).

The assessed morphological properties are presented in Figure 7. The height, width, and depth of dendritic arbors, including cell somas, was similar between pig and humans ICNs. While the number of dendrite stems, bifurcations, branches, and tips did not differ between pig and human cells, the total dendrite length (949 ± 571 µm vs 2009 ± 739 µm, P = 0.0025) as well as total dendrite surface area (5895 ± 4476 µm^2^ vs 17629 ± 12131 µm^2^, P = 0.023) were significantly greater at human ICNs. The greatest contributor to this difference appeared to be the number of short dendritic projections nearest the cell body (Fig 7K). Mouse ICNs which mostly (6/7) lacked dendrites were obviously more compact in size and, in the one cell with dendrites, projections were very short.

**Figure 7.**
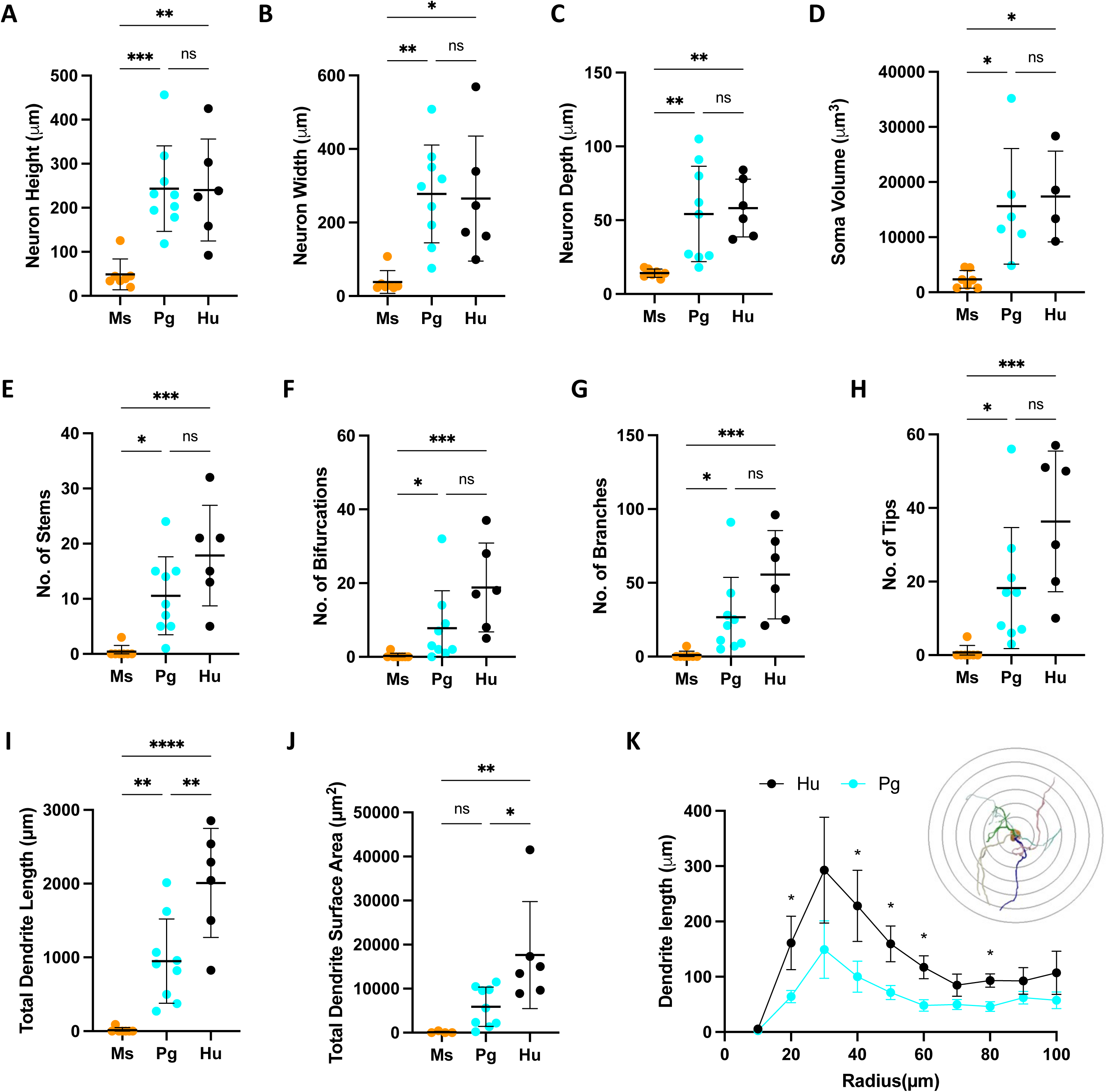
Quantification of morphological properties of mouse, pig and human ICNs. (A) Height, (B) width and (C) depth of neuron soma and dendrites. Mouse ICNs have smaller cell bodies and only 1/7 had dendrites. (D) The volume of the mouse ICN soma was significantly less than pig and human cells, which were similar in size. The number of dendritic stems (i.e. originating branch segments)(E), dendrite bifurcations (F), dendrite branches (G), and number of dendrite tips(H) were similar for pig and human cells, which were both greater than mouse ICNs. Human neuronal dendrites showed greatest total length (I) and total dendrite surface area (J). Sholl analysis (K) revealed the difference in dendrite length between pig and human neurons was greatest close to the cell body.

Across the small population of segmented mouse, pig and human ICNs, some variability in neuronal morphology was observed. In mouse, we found that not all axons were fully matured and showed growth cones at the distal end of the axon (Fig 6A_1_). From most cells we observed only a single axon according to the classification given above. In pig, a couple of cells appeared to lack any axon (Fig 6B_2_), and there was one human cell in which an axon could not be identified. In human, two cells lacked long dendritic projections and had only short dendritic projections constrained within an envelope around the soma (Fig 6C_2_). Every effort was made to completely delineate neuronal morphology, although, truncation due to either incomplete filling or imaging cannot be excluded. Whether these unique morphologies relate to a difference in function have not been determined. Also, whether the long intraganglionic projections were definitively dendrites or might possibly be collateral axons could not be determined at this time.

### ICN ultrastructure

Ultrastructural organization of the intracardiac ganglia increased with complexity from mouse to human. The differences observed exceed a simple increase in cell size but include an increased complexity of cytoplasmic organelles and a greater partitioning of the membrane in human and pig epicardial neurons. At mouse ICNs, synaptic specializations formed upon the somatic membrane forming axosomatic connections (Fig 8A,B). The synaptic terminals were filled with small clear vesicles (SCVs) and mitochondria. However, a few large dense core vesicles (LDCVs) were also observed among the SCVs (Fig 8B2). The mouse did not show any dendritic projections from the ICN somata, validating the results from light microscopy. In stark contrast, at pig ICNs, no axosomatic specializations were identified (Fig 8C). Instead, multiple protrusions of the somatic cytoplasm into dendritic processes were observed. Synaptic densities were challenging to locate but could be found at appositions between axonal and dendritic segments in areas adjacent to the neuronal cell bodies (Fig 8C2-4). These terminals were filled with mitochondria, SCVs, and some LDCVs.

**Figure 8.**
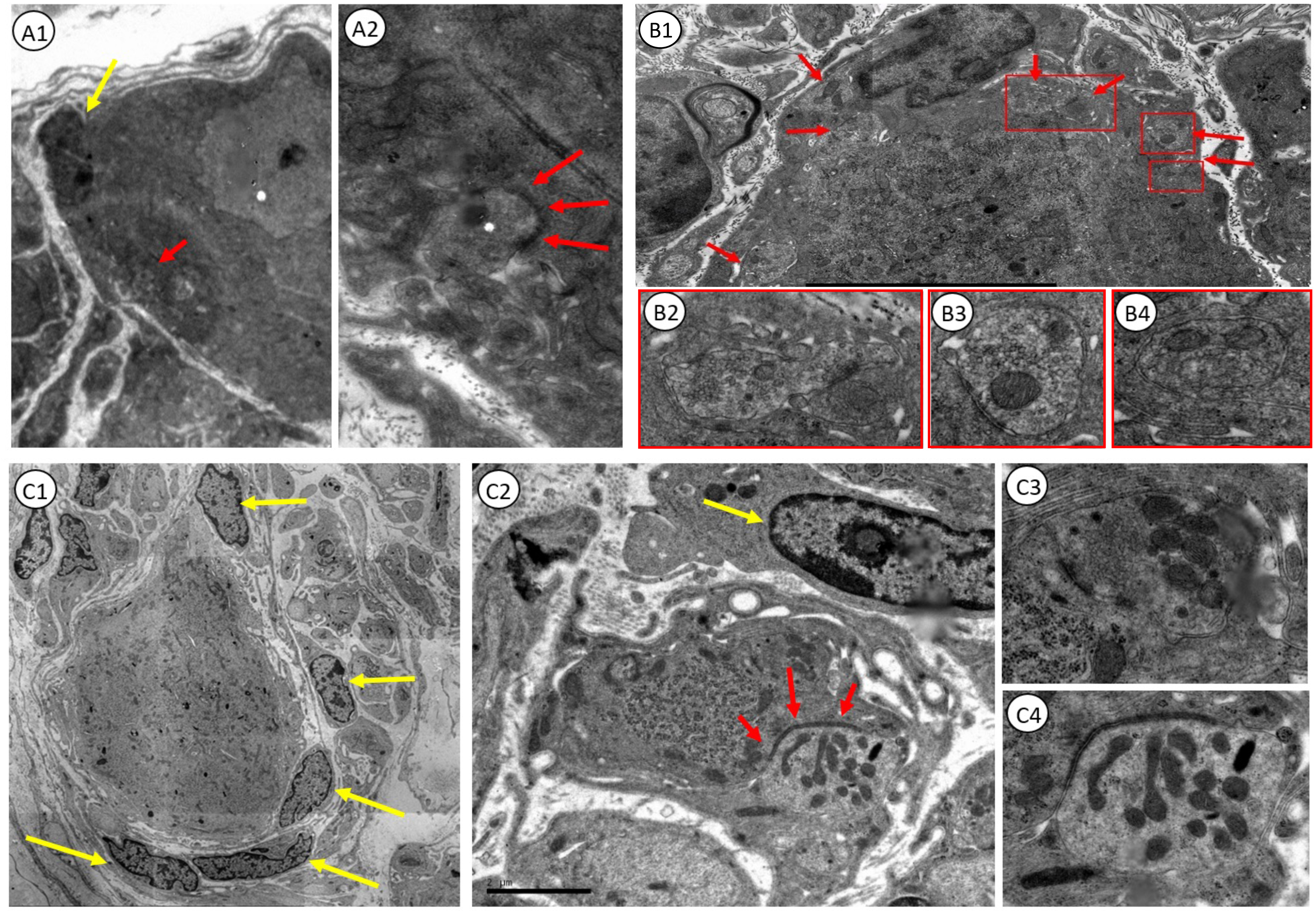
Ultrastructure of mouse and pig ICNs. TEM photomicrographs show ultrastructural features of mouse and pig ICNs. (A1) TEM image of a ganglionic neuron from a mouse epicardial ganglion. Yellow arrow indicates a satellite cell nuclei. Red arrow indicates a synaptic specialization forming on the somatic membrane. (A2) The structure indicated with red arrow in A1 is shown at higher magnification. Red arrows here indicate synaptic specialization on the somatic membrane of the mouse ICN. (B1) Another mouse ICN TEM image shows a satellite cell nucleus (*yellow arrow*) and multiple synaptic specializations on the somatic membrane (*red arrows*). Red boxed areas are expanded in B2, B3, and B4, which show ultrastructure of the axon terminal including small clear vesicles (SCVs) and mitochondria. (C1) TEM image of a ganglionic neuron from a pig intracardiac ganglion. Yellow arrows indica satellite cell nuclei. (C2) Synapse specialization forming on a dendrite in an area adjacent to the pig ICN soma. Synaptic specialization indicated with red arrows. Satellite cell nuclei indicated with yellow arrow. (C3, C4) Higher magnification images of axon terminals containing mitochondria and SCVs. C4 is the terminal shown in C2.

TEM of the somata of human ICNs typically revealed a nucleus with a prominent nucleolus and surrounding cytoplasm filled with varied quantities of vacuoles and lipofuscin granules (Fig 9). Some of the somata showed dendritic processes (Fig 9D). As in pig, axosomatic synapses were not encountered in the human ICN samples. Instead, axodendritic synaptic connections were present (Fig 9E). These axon terminals contained mitochondria, SCVs, and some LDCVs; and showed synaptic specialization with an electron dense active zone on the pre-terminal side of the synaptic cleft (Fig 9 E2), similar to axodendritic terminals in pigs.

**Figure 9.**
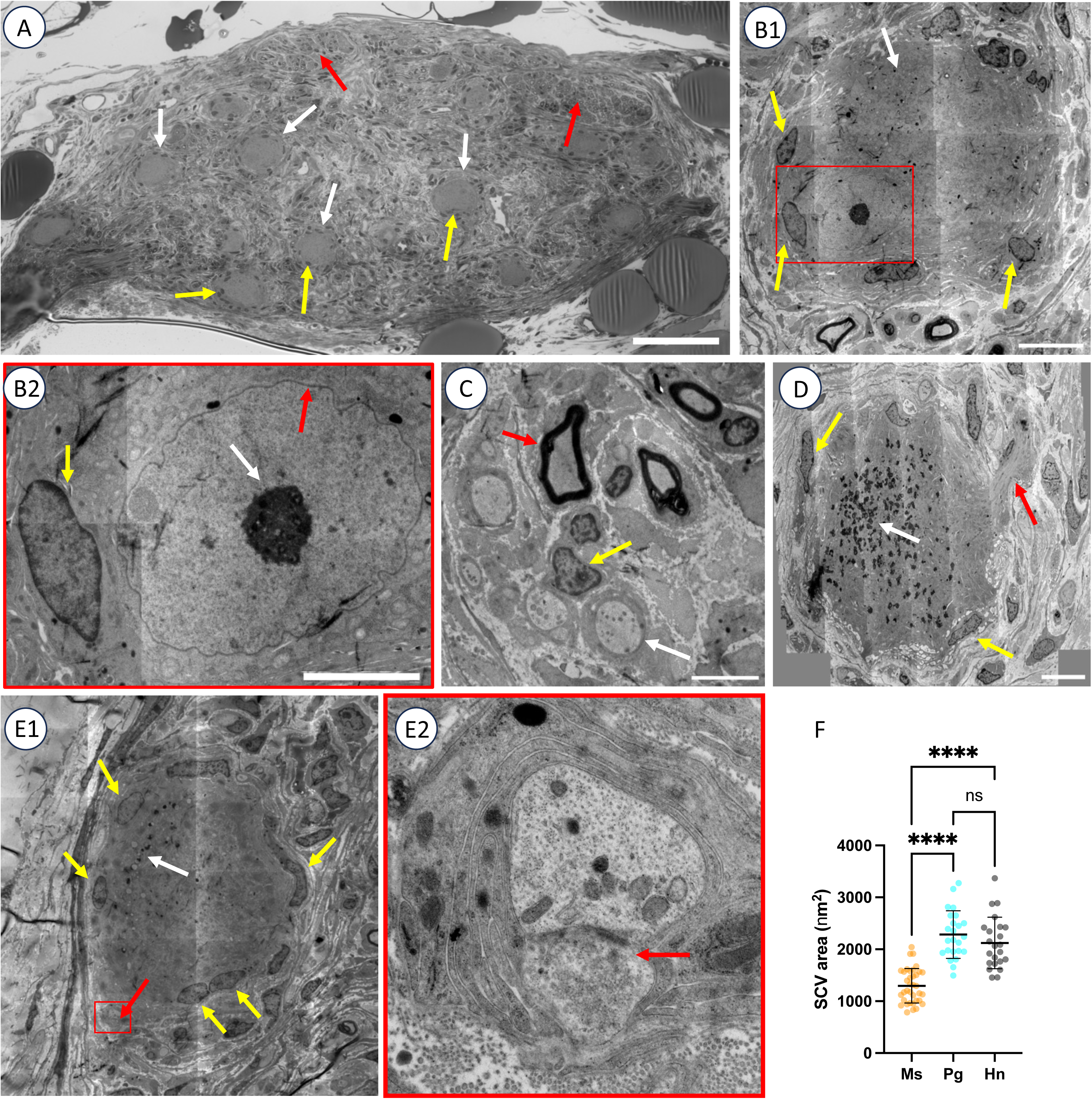
Ultrastructure of human ICNs. (A) Plastic embedded and toluidine blue section of human ICN organization. White arrows indicate neuronal somata. Yellow arrows indicate satellite cell nuclei. Red arrows indicate neuropil areas at the periphery of the ganglion with myelinated nerve fibers. (B1) TEM image of ganglionic neuron from human intracardiac ganglion. White arrow indicates cytoplasm of neuron. Yellow arrows show satellite cell nuclei. Red boxed area indicates position of nucleus at the periphery of the cell body. (B2) Higher magnification of boxed area in B1 with red arrow, white arrow, and yellow arrow indicating nuclear membrane, nucleolus, and satellite cell nucleus, respectively. (C) TEM image of neuropil area with red, yellow, and white arrows indicating a myelinated axon, Schwann cell nucleus, and unmyelinated axon. (D) TEM image of somata of human ICN with yellow, white, and red arrows indicating satellite cell nuclei, lipofuscin, and a dendritic process, respectively. (E1) TEM image of human ganglionic neuron with yellow, white, and red arrows indicating satellite cell nuclei, vacuoles and lipofuscin, and axodendritic synapse, respectively. (E2) Larger magnification of the red boxed area in E1 showing axodendritic synapse with SCVs, a few LDCVs, and a synaptic cleft with a pre-terminal active zone. The axon terminal is indicated with a red arrow. (F) Quantitative studies of SCV size between species. Note statistically larger porcine and human SCVs compared to SCVs in mice. There was no size difference between the porcine and human SCVs.

The cross-sectional areas of the SCVs in ICNs of mice, pigs, and humans were quantified from high resolution TEM images. All tissues had been processed for ultrastructural studies using identical fixation and staining protocols. The mean area of mouse SCVs (1,297 ± 331nm^2^, n = 36) was significantly smaller than SCVs at minipig (2283 ± 457 nm^2^, n = 24; P < 0.0001) and human ICNs (2121 ± 496 nm^2^, n = 23; P < 0.0001). (Fig 9 F). There was no difference in the size of minipig or human SCVs.

### Nerve evoked potentials at intracardiac neurons

Brief (100 μs) stimulation of the ganglion nerves elicited orthodromic and antidromic potentials at ICNs from mice, pigs, and humans (Fig 10). Single stimuli elicited fast, excitatory potentials which, at mouse ICNs, were always suprathreshold for the generation of an action potential. These cells exhibited a large excitatory post synaptic potential (EPSP)(Fig 10A). At pig and human ICNs, both subthreshold and suprathreshold EPSPs were observed. At all inputs tested with the ganglion nicotinic blocker hexamethonium (500uM), the potentials were blocked, almost completely, confirming activation of ganglion nicotinic receptors with acetylcholine as the principal mechanism for fast excitatory neurotransmission (data not shown). Antidromic potentials could be discerned from synaptic potentials in that they were all-or-none and were unaffected by hexamethonium.

**Figure 10.**
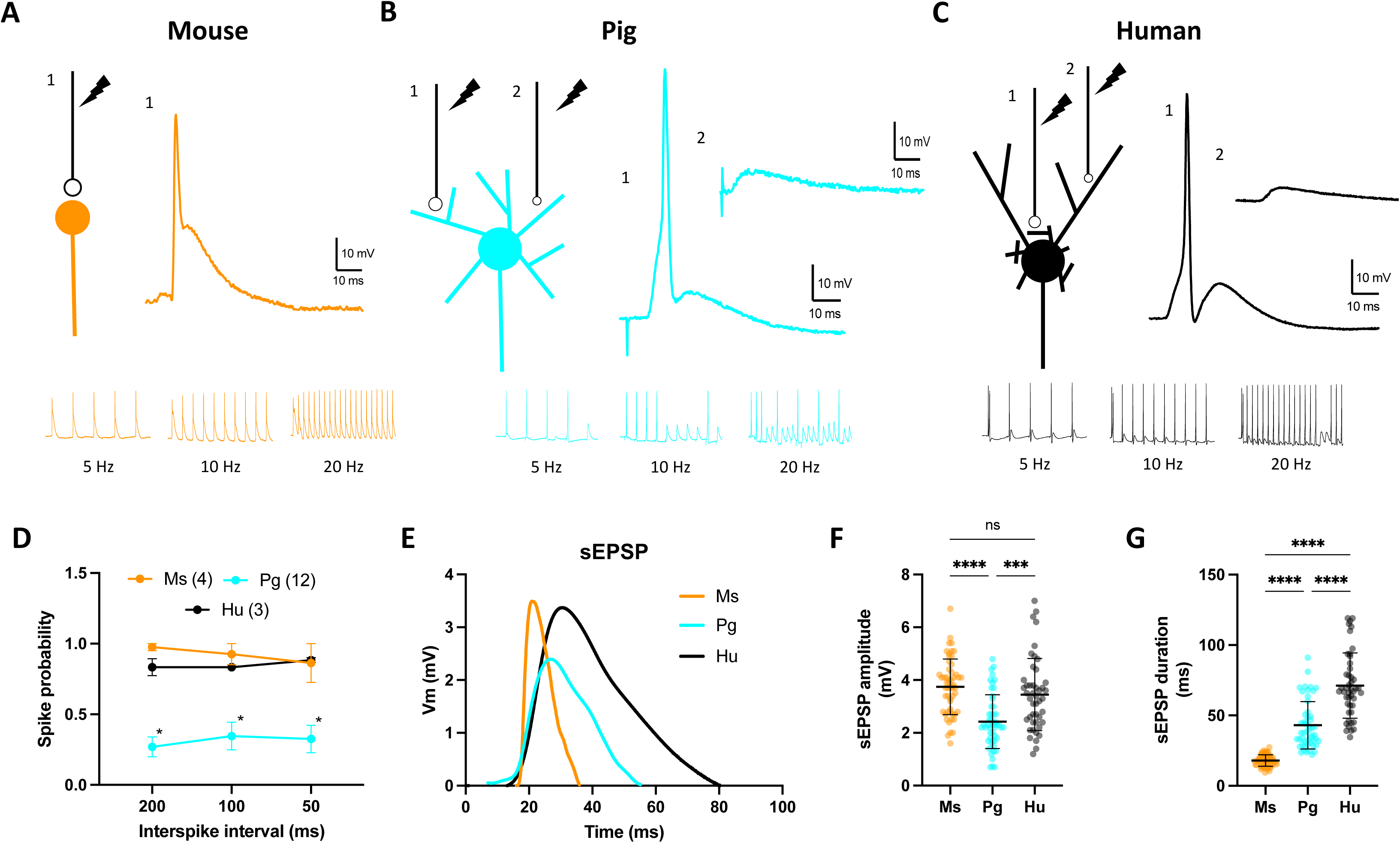
Assessment of ganglionic neurotransmission between species. Stimulation of preganglionic nerves evoked fast excitatory synaptic potentials (EPSPs) at mouse (A), pig (B) and human (C) ICNs. Representative responses are shown adjacent to a schematic illustration for synaptic connectivity as determined by light and electron microscopy. In mouse, single stimuli evoked suprathreshold EPSPs, while at pig and human ICNs, graded stimulation evoked both supra- and subthreshold EPSPs, providing evidence for convergence of cholinergic inputs. Synaptic potentials were blocked by the ganglion nicotinic antagonist hexamethonium (500 µM, data not shown). Shown below these traces are representative trains of stimuli, tested at multiple frequencies, which evoke synaptic potentials. (D) The probability of a stimulation evoked action potential is shown for each spike interval. Multiple failures for spike generation at pig ICNs were observed resulting in low synaptic efficacy for the tested synapses. (E) Spontaneous EPSPs (sEPSPs) were also observed at mouse, pig and human ICNs. Shown are representative tracings for an average sEPSP from mouse, pig and human ICN. Note that pig sEPSPs were significantly smaller than mouse or human potentials (F), while mouse sEPSPs had shortest duration and the spontaneous mini potentials at human ICNs had the longest duration.

To determine if the ganglionic neurons received multiple inputs, we used either graded stimuli at a single input, or, where possible, multiple ganglionic inputs were stimulated with independent extracellular electrodes. At pig ICNs, stimulation of ganglionic nerves evoked fast synaptic potentials. Most of the nerve evoked responses (32/35) were orthodromic, with only three antidromic spikes. 8 of 28 synapses tested received two discrete inputs based on latency alone. 9 of 15 cells tested for dual inputs with multiple electrodes had them. At human ICNs, nerve evoked potentials were obtained in 7/17 ganglion preparations. 5/7 of the potentials were identified as antidromic. In 3/7 preparations, the synaptically mediated EPSPs were graded in nature, showed variability in synaptic latency, and were sensitive to inhibition by hexamethonium (500 μM). Similar to mouse, a large suprathreshold EPSP was observed, which occasionally, due to the excitability of the post-synaptic membrane, elicited multiple spikes (Fig 10C). In one cell, two synaptically-evoked responses were observed after stimulation of a single nerve, with unique latency values that were both inhibited by hexamethonium, supporting the concept of convergent innervation, as observed for pig but not mouse ICNs.

In addition to single stimuli, trains of stimuli (5, 10, and 20 Hz) were given to test for presence of slow potentials and to measure synaptic efficacy. At pig ICNs, of 18 cells tested for slow potentials, 8/18 exhibited a slow hyperpolarization. No slow depolarizations were observed out of the 18 cells tested. Synaptic efficacy tended to be very low at porcine ICNs, with fewer than 40% of the evoked potentials triggering spikes. In contrast, at mouse ICNs, the evoked potentials elicited spikes nearly 100% of the time, and at human ICNs multiple spikes were observed with each depolarization giving greater than 100% synaptic efficacy (Fig 10C,D).

Spontaneous EPSPs (sEPSPs) were observed in mouse, pig and human ICNs. sEPSPs are mediated by spontaneous vesicle fusion, release of transmitter, and quantal-like activation of post-synaptic receptors. Representative sEPSPs from mouse, pig and human ICNs are shown in Figure 10E. Significantly smaller sEPSPs were observed at pig ICNs (2.3 ± 0.9 mV, n = 103), in comparison to mouse (4.5 ± 2.2 mV, n = 136; P < 0.0001) or human (4.4 ± 0.6 mV, n = 22; P < 0.0001), which were equivalent in size. The duration, measured from depolarization initiation to return to resting membrane potential, of mouse sEPSPs (18 ± 4.1 ms) was significantly shorter than those from pig (43 ± 17 ms; P < 0.0001) or human (71 ± 23 ms; P < 0.0001), and the human sEPSPs lasted longer than pig (P < 0.0001).

## Discussion

We identified structural and functional specification of human ICNs relative to two commonly used laboratory mammals, mice and pigs. Mice offer many advantages given available molecular tools and genetic manipulation, while pigs offer a preclinical model with potential for cardiac xenotransplantation. Across species we identify: 1) a scaling of somatic innervation density inversely proportional to dendritic arborization, 2) unique species dependent profiles of ganglionic neuropeptide expression, and 3) unique regulation of intrinsic membrane excitability. These properties shape intracardiac neuronal processing which by extrapolation would be distinct among these species. Human ICNs share structural similarity with the larger ICNs from pigs but exhibit membrane properties and neuropeptide expression patterns more common to mouse. The human cells showed hyperexcitable membranes, perineuronal sympathetic fibers, presence of VIP, the largest AHP amplitudes, greater proximal dendrite density, and the largest spontaneous synaptic potentials.

Changes in cell size and synaptic complexity among species has been shown for sympathetic^40^ and parasympathetic neurons^41^. Here we identify consistency with these scaling paradigms which now include data from human ICNs. The compact and densely organized ICNs of mice, have few if any dendrites observed with both light and electron microscopy. Accordingly, the density of axosomatic synapses increased. A reduced convergence of axonal inputs was also identified. This is consistent with the finding of greater somatic innervation at some sympathetic neurons which have reduced dendritic area^42^. The larger ICNs of pigs and humans grow elaborate dendritic arbors where convergence of axonal input was observed. We cannot say for certain that all long neurite projections were in fact dendrites or whether these may function as collateral axons to nearby cells. Interestingly both ‘dendrites’ and axons appeared to originate from adjacent membrane locations. We also observed that axons from pigs and humans ICNs often bifurcated near the soma giving rise to short axonal collaterals. We cannot yet determine if these are functional connections to adjacent cells. Given the unique morphology among large mammals, factors governing dendrite growth and maturation should prove informative to understanding factors regulating synaptic integration at the human heart.

Three classes of ICNs based on neurotransmitter contents are classically identified including cholinergic, adrenergic, and nitrergic cells (*for review see* ^5^). The most abundant cell type in each species is cholinergic neurons, identified in this study by the presence of the vesicular acetylcholine transporter (VAChT). Neurons with immunoreactivity for TH are also identified, yet, from previous observations, not all TH+ cells are believed to be functionally adrenergic, particularly in mice, due to the absence of proteins required for further synthesis and release of norepinephrine^43^. TH-IR varicosities have not been identified surrounding mouse ICNs, indicating a limited convergence of sympathetic terminals onto intracardiac ganglia in this species^44^. Based on previous immunohistochemical studies, functionally adrenergic neurons are thought to exist within human intracardiac ganglia^45^. We clearly identify TH-IR varicosities surrounding human ICNs, indicating the likely convergence of sympathetic nerves onto postganglionic neurons within human epicardial ganglia. We identified very few nitrergic cells or fibers within ICNs of mice, pigs, or humans, though they were previously observed in human RAGP removed during bypass surgery^32^, and in guinea pigs^46,47^. Remodeling of neurochemical expression in cardiac disease requires further study. These new data support unique functional roles of these classical fast neurotransmitters across species.

The expression and localization of neuropeptides was also unique between mice, pigs, and humans. The sensory neuropeptides SP and CGRP are found in some afferent neurons of DRG and nodose ganglia. SP causes slow depolarization at ICNs of rat, rabbit, and guinea pigs^48–52^. Interganglionic SP/CGRP fibers were identified in all three species, though perineuronal varicosities were only observed in mouse and human ganglia. The apparent density of SP/CGRP fibers was also noticeably lacking in pig intracardiac ganglia. The peptides termed NPY and VIP are commonly identified within sympathetic^53,54^ and parasympathetic^35,55^ neurons, respectively, and can modulate ICN activity^34,56–59^. A network of nerve fibers immunoreactive for NPY were abundant in human ganglia, but lesser so in ganglia from mice or pigs. NPY+ ICNs were identified in all three species suggesting that NPY might be co-released from intracardiac neurons within the heart. Ganglion fibers immunoreactive for VIP were also much more prevalent within human ganglia, with little to none observed in pigs and only a few VIP+ fibers observed in mice. Importantly, the presence of the peptide does not, in and of itself, demonstrate a functional role of these molecules. Additional work is necessary to confirm the unique expression profiles, localization, and activities of these peptides within human epicardial ganglia.

Like all neurons, the excitability of ICNs is dictated by the expression, localization, and activity of ion channels in the plasma membrane. Much work has identified roles for ionic conductances in the regulation of intrinsic excitability of ICNs from mice^60^, rats^37,38,61–66^, guinea pigs^52,67,68^, pigs^22,69–71^, and dogs^72–76;^ however, no data exists for human ICNs. Neuronal excitability is regulated in part by the non-inactivating voltage-dependent K^+^ current I_M_, the hyperpolarization-activated cationic conductance I_h_, and the transient outward K^+^ current I ^38^ Hyperpolarizing membrane currents identified the characteristic ‘sag’ indicative of activation of I_h_ in all three species. This first identification of H-current in human ICNs is clinically relevant due to prescribed use of the H-current blocker ivabradine for heart failure^77^, where it inhibits I_h_, also known as I_f_, at cardiac myocytes^78^. Ivabradine would likely have an effect at human ICNs. Additional work is necessary to identify the HCN channel subtypes expressed. Inhibition of A-current using 4-AP had no effect on human ICN excitability, moderately increased pig ICN excitability, and greatly increased the excitability of the mouse cells. Inhibition of SK channels with apamin greatly increased the number of spikes triggered by depolarization, identifying an important functional role for the small-conductance Ca^2+^-activated K^+^ current in regulation of membrane excitability of all three species. Following inhibition of SK channels, mouse neurons reached a maximum firing rate of 60hz, while pig and human neurons peaked at 20Hz and 26Hz, respectively. This suggests, that while the general ionic conductances are conserved, the kinetics of these channels may be uniquely tuned within each species, with mouse ICNs capable of the highest frequency of nerve activity. This higher frequency of nerve activity is likely associated with the higher intrinsic heart rate and greater need for high frequency vagal firing in mice.

Electron microscopy identified both similarities and differences in the ultrastructural organization of ICNs between species. The present samples of synaptic terminals in the ICN of mouse, pig, and human tissues show predominantly SCVs with presence of a few LDCVs, similar to previous reports based on TEM studies in these species^22,79,80^. However, the synaptic terminals are situated on the ICN somata in mice, whereas pigs and humans show axodendritic synapses. Similar axodendritic synapses have previously been reported in human ICNs^79^. A close relationship was also demonstrated across the studied species between the ICNs and glial processes. Similar to prior studies, satellite cell nuclei were readily identified around ICN somata and their glial processes extending to form near apposition with the ICN cell membranes^22,79^. Similar glial processes were also in close proximity to axodendritic synaptic contacts in pigs and humans. Although the synaptic terminals in ICNs shared many ultrastructural features between species, studies here of SCV size showed significantly larger SCVs in pigs and humans compared to mice, indicating species differences in transmitter quantities or packaging. These differences were supported by differences in the measured amplitudes of spontaneous postsynaptic potentials. Although synaptic contact is likely formed at higher resistance dendrites, these quantal-like events were significantly larger at ICNs from pig and human.

In summary, the importance of the intrinsic cardiac nervous system and its cellular components is well recognized and is increasingly being targeted in the management of human cardiac disease including atrial fibrillation, ventricular arrythmias, and heart failure^13–18,81,82^. Post-synaptic autonomic receptors at the heart have been the mainstay of cardiovascular pharmacology for the last half century. Increasingly more targeted approaches to manipulate the upstream neural circuitry, including the ICNs^83,84^, preganglionic fibers^19^, or afferent fibers^85^, are in clinical and preclinical testing. The use of human tissues is important to ensure that the findings obtained in research animals can be generalizable to human systems. The identification of ICN specialization across species described herein identifies areas for greater need regarding the understanding of dendrite and synapse formation, the ionic currents regulating membrane excitability, and the actions of cardiac neuropeptides unique to humans. While we are just beginning to scratch the surface of the many subtleties at the cellular level of human ICNs, we have made a significant step forward in the understanding of the uniqueness of these cells among species.

## Acknowledgements

We are grateful to those who have generously opted to donate their bodies, tissues, and organs posthumously, with the noble intention of saving lives and furthering biomedical research. Human tissue procurement was facilitated by Ms. Amiksha S. Gandhi. The image of the whole human heart shown in Figure 1C_1_ was made by Dr. Shumpei Mori. Surgical assistance with collection of porcine hearts was provided by Drs. Michael Dacey, Peter Hanna, and Joseph Hadaya. Ms. Amit Tsanhani assisted with porcine neuron staining, imaging, and morphological analysis. TEM samples were processed within the UCLA Brain Research Institute, Microscopic Techniques Laboratory by Dr. Chunni Zhu.

## Author contributions

{John D. Tompkins} Design, direction, and coordination of all work. Isolation of ganglia for IHC and TEM. Performed intracellular microelectrode recording, ganglion whole mount IHC, and single cell morphometry. Wrote manuscript and created figures.

{Donald B. Hoover} Design and direction of sectioned tissue IHC work.

{Leif A. Havton} Design and direction of TEM work.

{Janaki Patel} Assisted JDT with analysis of electrophysiological data as well as single cell segmentation and morphological analysis.

{Youngjin Cho} Assisted JDT with analysis of porcine cell neurophysiology.

{Elizabeth H. Smith} Performed IHC and imaging for DBH.

{Natalia Biscola} Performed TEM imaging and analysis for LAH.

{Olujimi A. Ajijola} Managed procurement of human tissues, critical review of the manuscript.

{Kalyanam Shivkumar} Critical review of the manuscript, funding.

{Jeffrey L. Ardell} Critical review of the manuscript, funding.

## Abbreviations: (in order of appearance)

ICNs: Intrinsic Cardiac Neurons
VIP: Vasoactive Intestinal Polypeptide
SP: Substance P
CGRP: Calcitonin Gene-Related Peptide
SAN: Sinoatrial node
AV: Atrioventricular
RAGP: Right Atrial Ganglionated Plexus
SPARC: Stimulating Peripheral Activity to Relieve Conditions
PSS: Physiologic Salt Solution
SVC: Superior Vena Cava
IVC: Inferior Vena Cava
sEPSPs: Spontaneous Excitatory Post-Synaptic Potentials
PBS: Phosphate Buffered Saline
VAChT: Vesicular Acetylcholine Transporter
TH: Tyrosine Hydroxylase
IR: Immunoreactive
VMAT2: Vesicular Monoamine Transporter 2
nNOS: Neuronal Isoform of Nitric Oxide Synthase
NPY: Neuropeptide Y
AHP: After Hyper-polarization
EPSP: Excitatory Post-Synaptic Potentials
SCVs: Small Clear Vesicles
LDCVs: Large Dense Core Vesicles
DRG: Dorsal Root Ganglia
ICG: Intrinsic Cardiac Ganglia

---

**Supplemental Table 1.** Organ donor demographic, heart transport time, and ECG data.

| Age | Race | Sex | ECG Notes |
| --- | --- | --- | --- |
| 52 | Caucasian | F | Sinus tachycardia, low voltage QRS, cannot rule out anterior infarct, abnormal ECG |
| 57 | Caucasian | M | NA |
| 44 | Hispanic | M | Normal Sinus rhythm, ST & T wave abnormality, consider inferolateral ischemia, Prolonged QT, Abnormal ECG |
| 48 | Hispanic | M | Sinus tachycardia, ventricular premature complex, old inferior infarct |
| 72 | Caucasian | M | Atrial fibrillation with rapid ventricular response. Moderate ST depression. Abnormal rhythm ECG. |
| 57 | Hispanic | M | Normal heart rhythm, normal ECG |
| 64 | Asian | M | Normal sinus rhythm, T wave abnormality, consider anterior ischemia, abnormal ECG |
| 62 | African American | F | Normal sinus rhythm, prolonged QT, abnormal ECG, low voltage QRS, nonspecific T wave abnormality |
| 63 | Hispanic | M | Sinus rhythm with marked sinus arrhythmia, left anterior fascicular block, anterolateral infarct, age undetermined, abnormal ECG |
| 40 | Caucasian | M | Sinus rhythm, low voltage, precordial leads, abnormal T, consider ischemia, lateral leads, Abnormal ECG |
| 52 | Hispanic | M | Sinus tachycardia, Moderate T-wave abnormality, consider later and inferior ischemia, Abnormal ECG |
| 66 | Hispanic | F | Atrial fibrillation and slow ventricular response, abnormal ECG |
| 64 | Hispanic | M | Sinus tachycardia, T wave abnormality, consider inferior ischemia. Abnormal ECG |
| 66 | Hispanic | F | Sinus tachycardia, abnormal R-wave progression, early transition. Borderline T wave abnormality |
| 47 | Hispanic | M | NA |
| 57 | African American | M | Sinus rhythm with first degree AV block, Low QRS voltage, Prolonged QT interval, Abnormal ECG |
| 48 | Caucasian | F | Sinus rhythm, long QT, abnormal ECG |

